# ELMOD1 stimulates ARF6-GTP hydrolysis to stabilize apical structures in developing vestibular hair cells

**DOI:** 10.1101/189621

**Authors:** Jocelyn F. Krey, Rachel A. Dumont, Philip A. Wilmarth, Larry L. David, Kenneth R. Johnson, Peter G. Barr-Gillespie

## Abstract

Sensory hair cells require control of physical properties of their apical plasma membranes for normal development and function. Members of the ARF small GTPase family regulate membrane trafficking and cytoskeletal assembly in many cells. We identified ELMOD1, a guanine nucleoside triphosphatase activating protein (GAP) for ARF6, as the most highly enriched ARF regulator in hair cells. To characterize ELMOD1 control of trafficking, we used a mouse strain lacking functional ELMOD1 (roundabout; rda). In rda/rda mice, cuticular plates of utricle hair cells initially formed normally, then degenerated after postnatal day 5 (P5); large numbers of vesicles invaded the compromised cuticular plate. Hair bundles initially developed normally, but the cell’s apical membrane lifted away from the cuticular plate, and stereocilia elongated and fused. Membrane trafficking in type I hair cells, measured by FM1-43 dye labeling, was altered in rda/rda mice. Consistent with the proposed GAP role for ELMOD1, the ARF6 GTP/GDP ratio was significantly elevated in rda/rda utricles as compared to controls, and the level of ARF6-GTP was correlated with the severity of the rda/rda phenotype. These results suggest that conversion of ARF6 to its GDP-bound form is necessary for final stabilization of the hair bundle.

**Significance Statement:** Assembly of the mechanically sensitive hair bundle of sensory hair cells requires growth and reorganization of apical actin and membrane structures. Hair bundles and apical membranes in mice with mutations in the *Elmod1* gene degenerate after formation, suggesting that the ELMOD1 protein stabilizes these structures. We show that ELMOD1 is a GTPase-activating protein in hair cells for the small GTP-binding protein ARF6, known to participate in actin assembly and membrane trafficking. We propose that conversion of ARF6 into the GDP-bound form in the apical domain of hair cells is essential for stabilizing apical actin structures like the hair bundle and ensuring that the apical membrane forms appropriately around the stereocilia.

## Introduction

A hair cell in the inner ear transduces auditory and vestibular stimuli with its hair bundle, a mechanically sensitive cluster of actin-rich stereocilia that emanate from the cell’s apex (Gillespie and Müller, 2009). Apical membranes play a unique, fundamental role in inner-ear function and, accordingly, apical and basolateral membranes of hair cells differ in their phospholipid and membrane-protein content (Zhao et al., 2012). Hair cells have been reported to recycle their apical membranes at a rate of >1 μm^2^/s, leading to a complete turnover of the apical membrane in less than 1 min (Griesinger et al., 2002; Griesinger et al., 2004). Moreover, membrane tension likely regulates stereocilia length (Prost et al., 2007; Manor and Kachar, 2008) and mechanotransduction (Powers et al., 2012; Powers et al., 2014; Peng et al., 2016). Control of the balance between exo‐ and endocytosis is critical, as a slight mismatch can lead to morphological abnormalities like membrane blebbing (Shi et al., 2005).

The ADP-ribosylation factor (ARF) family of small GTPases, which includes the classic ARFs and the ARF-like (ARL) proteins, regulates membrane trafficking in most cell types (D’Souza-Schorey and Chavrier, 2006). ARF6 in particular regulates endosomal membrane traffic and apical membrane structure, through both clathrin-mediated and clathrin-independent apical endocytosis (Doherty and McMahon, 2009). ARF6 also regulates cellular actin through its interactions with RHO family members like RAC1 and CDC42 (Franco et al., 1999; Radhakrishna et al., 1999; Hyman et al., 2006; Klein et al., 2006; Osmani et al., 2010). Like other small GTPases, the ARFs are activated by guanine-nucleotide exchange factors (GEFs) and inhibited by GTPase-activating proteins (GAPs). The ELMO domain containing protein family (ELMODs) consists of six paralogs in mammals, two of which (ELMOD1 and ELMOD2) have been shown to be ARF-family GAPs (East et al., 2012; Ivanova et al., 2014). ELMOD1 had its greatest GAP activity towards ARF6, while ELMOD2 was a GAP for ARL2 (Ivanova et al., 2014).

Mutations in ELMOD1 lead to deafness and vestibular dysfunction in mice (Johnson et al., 2012). The *roundabout* (*rda*) allele has a large deletion in the first half of the *Elmod1* gene, which results in loss of ELMOD1 protein expression (Johnson et al., 2012). Beginning after postnatal day 7 (P7), mice homozygous for the *rda* mutation (*rda/rda*) develop morphological abnormalities in cochlear hair cells, including fusion and elongation of inner hair cell stereocilia and degradation of outer hair cell stereocilia (Johnson et al., 2012). Little is known of the consequences of the *rda* mutation on vestibular hair cells, however, although circling exhibited by the homozygous mutant mice suggests a lack of vestibular function. ELMOD1 is also expressed within the brain, and has been detected in cerebellar Purkinje cells and granule cells and pyramidal neurons within the hippocampus (Johnson et al., 2012). Interestingly, a mutation in *Elmod3* was recently linked to deafness in humans as well (Jaworek et al., 2013), confirming the significance of this protein family for inner-ear function.

To examine the role of ELMOD1 in mouse vestibular hair cells, we determined by immunoblotting that it is developmentally regulated, peaking near the end of vestibular hair cell development. Hair cells initially developed normally in *rda/rda* mice, but by P5, defects in the cuticular plate were observed, followed by stereocilia degeneration. Like with mouse mutations in *Myo6*, *Ptprq*, and *Rdx*, apical membranes lifted up and the stereocilia actin cores elongated and fused, leaving in the most extreme cases a single giant stereocilium. We used *rda/rda* mice to demonstrate the GAP activity of ELMOD1 toward ARF6, suggesting that the consequences of the *Elmod1* mutation were due to elevated ARF6-GTP levels. We propose that ARF6 must be converted to the GDP form at apical surfaces to permit stabilization of the hair bundle’s actin and membrane structures.

## Materials and Methods

### Nomenclature

Per convention (http://www.informatics.jax.org/mgihome/nomen/gene.shtml), all protein names use the official gene symbol (http://www.genenames.org) with all caps and no italics.

### Mice

*Roundabout* (*rda*) mice on a C57BL/6J background were obtained from The Jackson Laboratory. To obtain mice heterozygous and homozygous for the *rda* mutation, +*/rda* females were crossed to *rda/rda* males. These crosses allowed us to generate equal numbers of knockout mice and heterozygote controls, which was especially important for proteomics experiments. C57BL/6 mice were used as wildtype controls (referred to as B6).

### Experimental design and statistical analyses

Because +*/rda* mice have normal auditory and vestibular function (Johnson et al., 2012), only comparisons of +*/rda* to *rda/rda*—which have the same genetic background—were functionally relevant. Because of the different genetic backgrounds, B6 comparisons to +*/rda* and *rda/rda* are of less relevance for determining mechanisms. The Student’s t-test was used for all pairwise comparisons (two-sided, two-sample equal variance). Data distribution was assumed to be normal but this was not formally tested.

### Mass spectrometry of TMT-labeled utricle extracts

Utricles were dissected from +*/rda* and *rda/rda* mice at P12. Four biological replicates were prepared for each genotype with 4-6 utricles per replicate. Lysates were prepared using the eFASP method (Erde et al., 2014). Briefly, lysis buffer (4% SDS, 0.2% deoxycholic acid) at 15 μl per utricle was added to each tube and samples were vortexed, heated to 90°C for 10 min, and bath sonicated for 5 min. Protein concentration of each lysate was measured using the Micro BCA Protein Assay Kit (ThermoFisher). Lysates were divided into 2 μg aliquots. Samples were then digested as described (Erde et al., 2014), with triethylammonium bicarbonate (TEAB) replacing the ammonium bicarbonate in all solutions.

Peptides were each reconstituted in 25 μl of 100 mM triethylammonium bicarbonate (TEAB) and labeled with 10-plex Tandem Mass Tag (TMT) reagents from Thermo Scientific. TMT reagents (0.8 g) were each dissolved in 52 μl anhydrous acetonitrile (ACN). Each sample, containing 2 μg of peptide in 25 μl volume of TEAB buffer, was combined with 12 μl of its respective TMT reagent and incubated for 1 h at room temperature. Two μl of each reaction mixture was then added, and the mixture was incubated at room temperature for 1 hr; 2 μl of 5% hydroxylamine was added, and the combined sample was incubated for a further 15 min. The mixture was dried down and dissolved in 5% formic acid. A 2 μg aliquot of labeled peptide was analyzed by a single 2 hr LC-MS/MS method using an Orbitrap Fusion as described below; this run was performed to normalize the total reporter ion intensity of each multiplexed sample and check labeling efficiency. The remaining samples were quenched by addition of 2 μl of 5% hydroxylamine as above, combined in a 1:1:1:1:1:1:1:1:1:1 ratio based on total reporter ion intensities determined during the normalization run, and dried down in preparation for 2D-LC-MS/MS analysis.

Multiplexed TMT-labeled samples were reconstituted in 5% formic acid and separated by twodimensional reverse-phase liquid chromatography using a Dionex NCS-3500RS UltiMate RSLCnano UPLC system. A 20 μl sample (10.4 μg) was injected onto a NanoEase 5 μm XBridge BEH130 C18 300 μm × 50 mm column (Waters) at 3 μl/min in a mobile phase containing 10 mM ammonium formate at pH 9. Peptides were eluted by sequential injection of 20 μl volumes of 14, 20, 24, 28, 35, and 90% acetonitrile in 10 mM ammonium formate (pH 9) at 3 μl/min flow rate. Eluted peptides were diluted with mobile phase containing 0.1% formic acid at 24 μl/min flow rate and delivered to an Acclaim PepMap 100 μm × 2 cm NanoViper C18 (5 μm) trap on a switching valve. After 10 min of loading, the trap column was switched in-line to a PepMap RSLC C18, 2 μm, 75 μm × 25 cm EasySpray column (Thermo Fisher). Peptides were then separated at low pH in the 2nd dimension using a 7.5-30% acetonitrile gradient over 90 min in mobile phase containing 0.1% formic acid at 300 nl/min flow rate. Each 2nd dimension LC run required 2 hr for separation and re-equilibration, so each 2D LC-MS/MS method required 12 hr for completion. Tandem mass spectrometry data was collected using an Orbitrap Fusion Tribrid instrument configured with an EasySpray NanoSource (Thermo Fisher). Survey scans were performed in the Orbitrap mass analyzer (resolution = 120,000), and data-dependent MS2 scans were performed in the linear ion trap using collision-induced dissociation (normalized collision energy = 35) following isolation with the instrument’s quadrupole. Reporter ion detection was performed in the Orbitrap mass analyzer (resolution = 60,000) using MS3 scans following synchronous precursor isolation of the top ten ions in the linear ion trap, and higher-energy collisional dissociation in the ion-routing multipole (normalized collision energy = 65).

Confident peptide identifications and reporter ion peak heights were obtained using Proteome Discoverer, (v1.4, Thermo Fisher). SEQUEST search parameters were: parent ion tolerance of 1.25 Da, fragment ion tolerance of 1.0 Da, tryptic cleavage with no more than two missed cleavages, variable oxidation of Met, and static modifications for alkylation and TMT reagents. A custom protein database with complete coverage and minimal peptide redundancy was used. Protein database details and description of the quantitative analysis are provided in ProteomeXchange submission below; TMT data processing and statistical testing has been previously described (Plubell et al., 2017).

The mass spectrometry proteomics data have been deposited to the ProteomeXchange Consortium via the PRIDE partner repository (Vizcaíno et al., 2016) with the dataset identifier PXD006164.

### Immunoblotting

Hair bundles and utricles were isolated from C57BL/6J, +*/rda* and *rda/rda* mice and then were subjected to SDS-PAGE and immunoblotting as described previously (Shin et al., 2013). Western blots were probed with a 1:1000 dilution of anti-ELMOD1 antibody (NBP1-85094, Novus Biologicals; RRID:AB_11005087), a 1:100 dilution of anti-ARF6 antibody (3A1, sc-7971, Santa Cruz Biotechnology; RRID:AB_2289810), or a 1:2000 dilution of anti-actin antibody (JLA20, Developmental Studies Hybridoma Bank; RRID:AB_528068).

### Measurement of ARF6-GTP binding

To measure ARF-GTP levels, we used the ARF6 Pull-down Activation Assay Biochem Kit (Cytoskeleton, Inc. #BK033-S), following the manufacturer’s instructions with several modifications. Utricles were isolated from C57BL/6J, *+/rda*, and *rda/rda* mice at P11-P12, and otolithic membranes were removed using an eyelash. Four to eight organs were dissected at a time from each genotype and kept on ice during the dissection process. Utricles were transferred to a tube in a small amount of dissection solution; the tube was gently spun in a microcentrifuge and most of the solution was removed. Lysis buffer containing 1:100 Protease Inhibitor Cocktail (included in kit) was added to organs (10 μl lysis buffer per utricle). Utricles were probe-sonicated with 3 × 5 sec pulses at 25% amplitude while on ice, then flash frozen in liquid nitrogen and stored at ‐80°C. Frozen samples were gently thawed on ice, then spun at 4°C in a microcentrifuge at max speed for 2 min. A small aliquot (4-6 μl) of the solution was saved as total lysate sample, and the remaining solution was transferred to a new tube and brought up to a 100 μl volume with additional lysis buffer. Lysates were mixed on a rotator for 1 hr at 4°C with 10 μl (10 μg) of GGA3 beads, then spun down at 4500 × g at 4°C for 2 min. The supernatant was removed and beads were washed twice, each with 600 μl of wash buffer followed by a spin at 4500 × g at 4°C for 2 mins. The wash buffer was removed from beads and 10 μl of 2× LDS sample buffer with reducing agent (Thermo Fisher) was added to each tube. An equal amount of 2× LDS sample buffer was also added to the total lysates. Tubes were incubated for 2 min at 95°C and then samples were separated on a 1 mm 12-well 12% Bis-Tris gel (Life Technologies) run with MES running buffer. Gels were transferred for 1 hr at 22 V at 4°C onto 0.2 μm PVDF membranes (Immobilon P^SQ^, Millipore). Protein immunoblots were probed as described above using anti-ARF6 and anti-actin antibodies.

### *In utero* electroporation

*Elmod1* was cloned from a mouse brain cDNA library and inserted into a GFP-C1 vector. Pregnant female mice (C57BL/6 × CD1 crosses) were anesthetized and up to eight embryos at E10.5 to E12.5 were injected in the otocyst with <200 nl of the GFP-ELMOD1 construct. Surgical procedures were performed as described previously (Ebrahim et al., 2016).

### Immunocytochemistry and imaging

Using 4% formaldehyde in PBS, utricles were fixed for 4-12 hr and cochleas were fixed for 0.5-1 hr. Organs were then rinsed in PBS, permeabilized for 10 min in 0.5% Triton X-100 in PBS, and blocked for 1-2 hr in 2% bovine serum albumin and 5% normal goat serum in PBS. For immunolocalization of ARF6 and other membrane compartment markers, organs were permeabilized and blocked for 1 hr in 5% normal goat serum and 0.2% saponin in PBS; this blocking solution was used in all subsequent antibody steps. Organs were incubated overnight at 4°C with primary antibodies diluted in blocking solution; a dilution of 1:250 was used for anti-ELMOD1 and 1:1000 for anti-acetylated tubulin (Sigma-Aldrich T7451; RRID:AB_609894). For ARF6 and other membrane markers, the following dilutions were used: 1:100 anti-ARF6, 1:250-activated ARF6 (NewEast Biosciences, #26918; RRID:AB_2629397); 1:250 anti-clathrin heavy chain (X22, MA1-065, ThermoFisher Scientific; RRID:AB_2083179), 1:250 anti-EEA1 (#3288, Cell Signaling Technology; RRID:AB_2096811), 1:250 anti-Rab11 (#5589, Cell Signaling Technology; RRID:AB_10693925), 1:250 anti-Rab5 (#3547, Cell Signaling Technology; RRID:AB_2300649), 1:250 anti-LAMP1 (H4A3, Developmental Studies Hybridoma Bank; RRID:AB_2296838). Organs were then rinsed 3× for 10 min each, followed by a 3-4 hr incubation in blocking solution with 1:1000 Alexa Fluor secondary antibodies and 0.4 U/ml Alexa Fluor 488 Phalloidin (Molecular Probes, Invitrogen). Similarly, utricles from electroporated ears were fixed and stained as above using an anti-GFP (DSHB-GFP-4C9, Developmental Studies Hybridoma Bank; RRID:AB_2617422) primary antibody and TRITC-phalloidin. Following secondary antibody incubation, organs were then rinsed 3× for 20 min each and mounted on slides in Vectashield (Vector Laboratories) using a single Secure-Seal spacer (8 wells, 0.12 mm deep, Invitrogen). Images were acquired on an Olympus FluoView FV1000 laser scanning confocal microscope system and AF10-ASW 3.0 acquisition software, using a 60× 1.42 NA Plan Apo objective with 3× zoom and 0.4 μm z-steps. ARF6 and ARG-GTP images were collected on a Zeiss LSM 710 using either a 63× 1.4 NA Plan-Apochromat or 25× 0.8 NA Plan-Neofluar objective. Fiji software was used to adjust intensities, generate reslice stacks, and prepare stack projections. To analyze ARF6 and ARF6-GTP levels, average z-projections centered around the cuticular plate were generated for two separate areas within each 25× stack. Regions of interest (ROIs) surrounding the cuticular plate (as visualized by phalloidin) were generated for 40-50 cells per image, and mean intensities were measured in each channel using the multi-measure function within the ROI Manager tool in Fiji. To correct for global intensity differences between samples imaged on different days, measurements were corrected for background levels within each image and normalized to the total phalloidin average. To analyze the correlation between ARF6-GTP level and phenotype, ROIs were generated around the cuticular plate region of 40 cells within each 63× stack and ARF6-GTP intensities were measured using Fiji. Measurements from two separate experiments with images from two utricles per condition were acquired. Each of the 160 cells was qualitatively scored on six different morphological phenotypes on a 1 (noticeable) to 2 (very noticeable) point scale. The phenotypes scored were as follows: gaps in cuticular plate actin, protrusion of apical surface, movement of the bundle towards the fonticulus, decrease in bundle size, increase in long stereocilia, and fusion of stereocilia. The cell type (type 1 vs. type 2) and whether the bundle was newly developed (based on bundle size and stereocilia thickness) was also noted.

### Transmission electron microscopy

Utricles from P5 or P12 +*/rda* or *rda/rda* mice were immersion-fixed in 1% glutaraldehyde, 1% OsO_4_, 0.1 M phosphate for 3 hr, dehydrated in a graded series of acetone, and embedded in Araldite (Electron Microscopy Sciences). The Araldite was cured in a 60°C oven for 48 hr. Sections of 90 nm were collected on Maxtaform 200-mesh Cu/Rh grids (Ted Pella, Inc.), were stained using 1% uranyl acetate (Electron Microscopy Sciences) and Reynold’s lead citrate, and were examined on a Philips CM100 transmission electron microscope.

### Endocytosis assay using FM1-43FX

Utricles were dissected on ice, and then transferred to ice-cold Hanks balanced salt solution (HBSS) containing 100 μM tubocurarine to block transduction channels. Tubocurarine was included in all subsequent solutions. Organs were incubated on ice for 1 min with 20 μM FM1-43FX, rinsed briefly in cold HBSS, then were transferred to HBSS at 37°C for up to 1 hr. Some organs were transferred immediately to cold 4% formaldehyde for 20 min rather than being transferred to 37°C. All organs were fixed on ice for 20 min in 4% formaldehyde, then washed three times in cold PBS. Organs were mounted and imaged as above. Images from at least 2 separate experiments with 1-2 utricles per time point were analyzed using Fiji. The morphology of the hair cell neck region was used to distinguish between type I and type II hair cells, and ROIs were generated surrounding the cuticular plate of ten type I hair cells per image. Measurements were then made using this ROI at intervals of five sections above and below the cuticular plate, marking measurements made at the bundle, cuticular plate, pre-nuclear, nuclear, and post-nuclear levels.

## Results

### ARF family and ARF modulator expression

To determine whether ELMOD1 plays a unique role in hair cells, we examined expression of ARFs and ARF modulators like ELMOD1 in hair cells. We analyzed data from experiments that used the *Pou4f3-Gfp* mouse line to purify hair cells from mouse cochlea and utricle, then carried out RNA-seq to measure transcript abundance (Shen et al., 2015). In utricle (Fig. 1A), of all ARFs and ARF modulators, only *Elmod1* was enriched 10-fold or greater in hair cells; in cochlea, only *Adap1* and *Elmod1* were enriched 10-fold or greater, with *Elmod1* at a 360-fold higher level (Fig. 1B). These experiments showed that *Elmod1* was consistently enriched in hair cells, more so than any other ARFs or ARF modulators.

**Figure 1.**
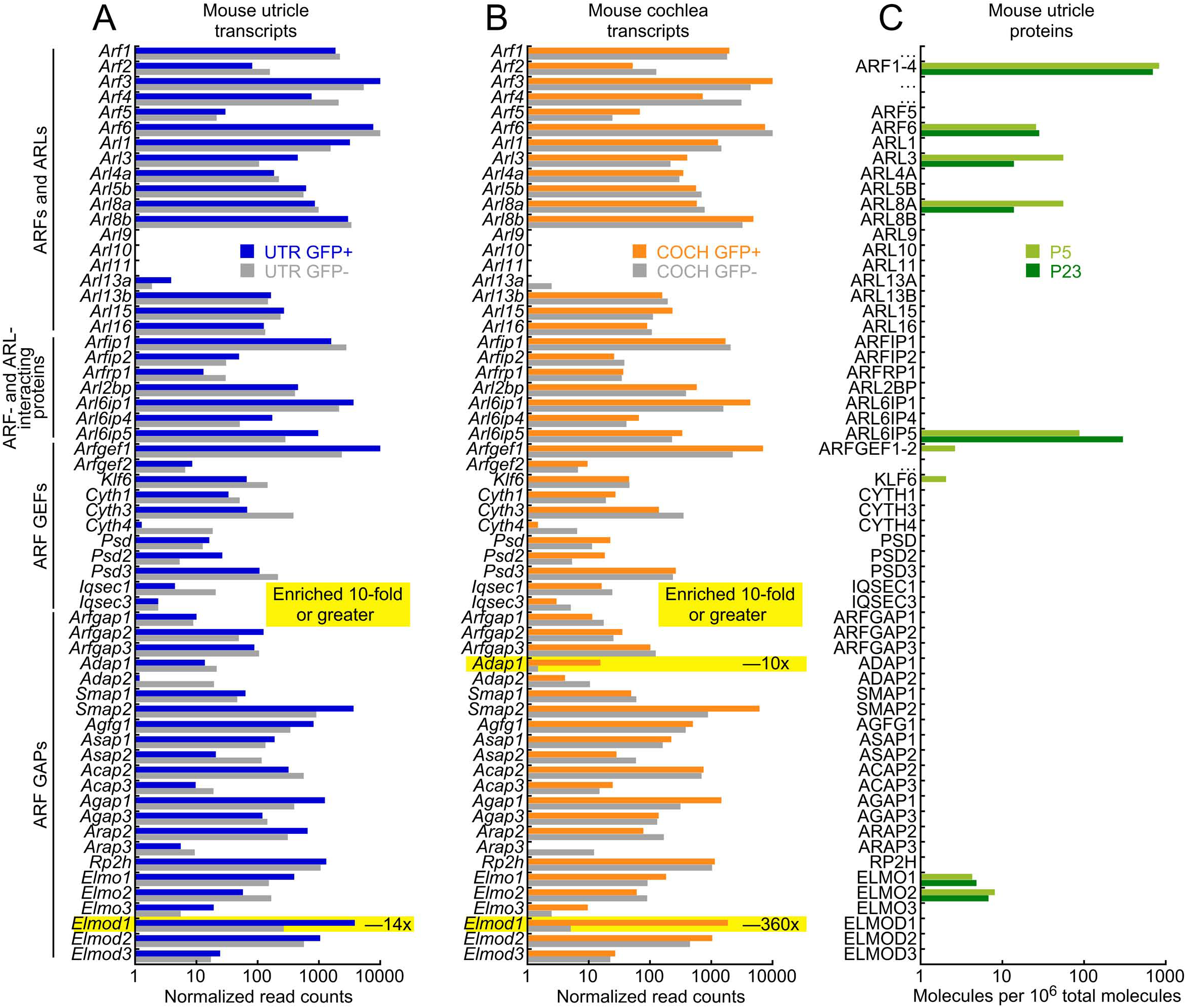
ARF and ARF modulator transcript analysis in hair cells. ***A***, Transcript levels in sorted hair cells (GFP+) and other inner ear cells (GFP-) from mouse utricle; data from Shen et al. (2015). ***B***, Cochlear transcript levels. ***C***, ARF and ARF-modulator proteins detected by mass spectrometry in P5 and P23 hair bundles and utricle; data from Krey et al. (2015). Because of shared peptides, ARFs 1-4 and ARFGEFs 1 and 2 could not be unambiguously quantified individually and thus were combined.

We also examined protein levels of ARFs and ARF regulators (Fig. 1C). In mouse utricle extracts (Krey et al., 2015), we detected the ARF family members ARF1-4 (combined because of ambiguity from shared peptides), ARF6, ARL3, and ARL8A. We detected several ARF modulators, but did not reliably observe ELMOD1 in mouse inner ear samples using mass spectrometry. That discrepancy with transcriptomics experiments likely reflects the difference in sensitivity and dynamic range between the two approaches.

### Expression of ELMOD1 in utricles

Although ELMOD1 was not detected by mass spectrometry, a sensitive antibody against ELMOD1 readily detected a band of the correct molecular mass (38 kD) in utricle hair bundles, whole utricle, and brain extracts (Fig. 2A-B). The 38 kD band disappeared in bundles or whole utricle from *rda/rda* mice (Fig. 2A). While several higher molecular mass bands persisted, the 38 kD band also disappeared completely in *rda/rda* brain extracts (Fig. 2B). ELMOD1 was expressed in the mouse utricle as early as embryonic day 18 (E18), and increased in levels relative to actin across postnatal development, with peak levels of expression at P7 (Fig. 2B).

**Figure 2.**
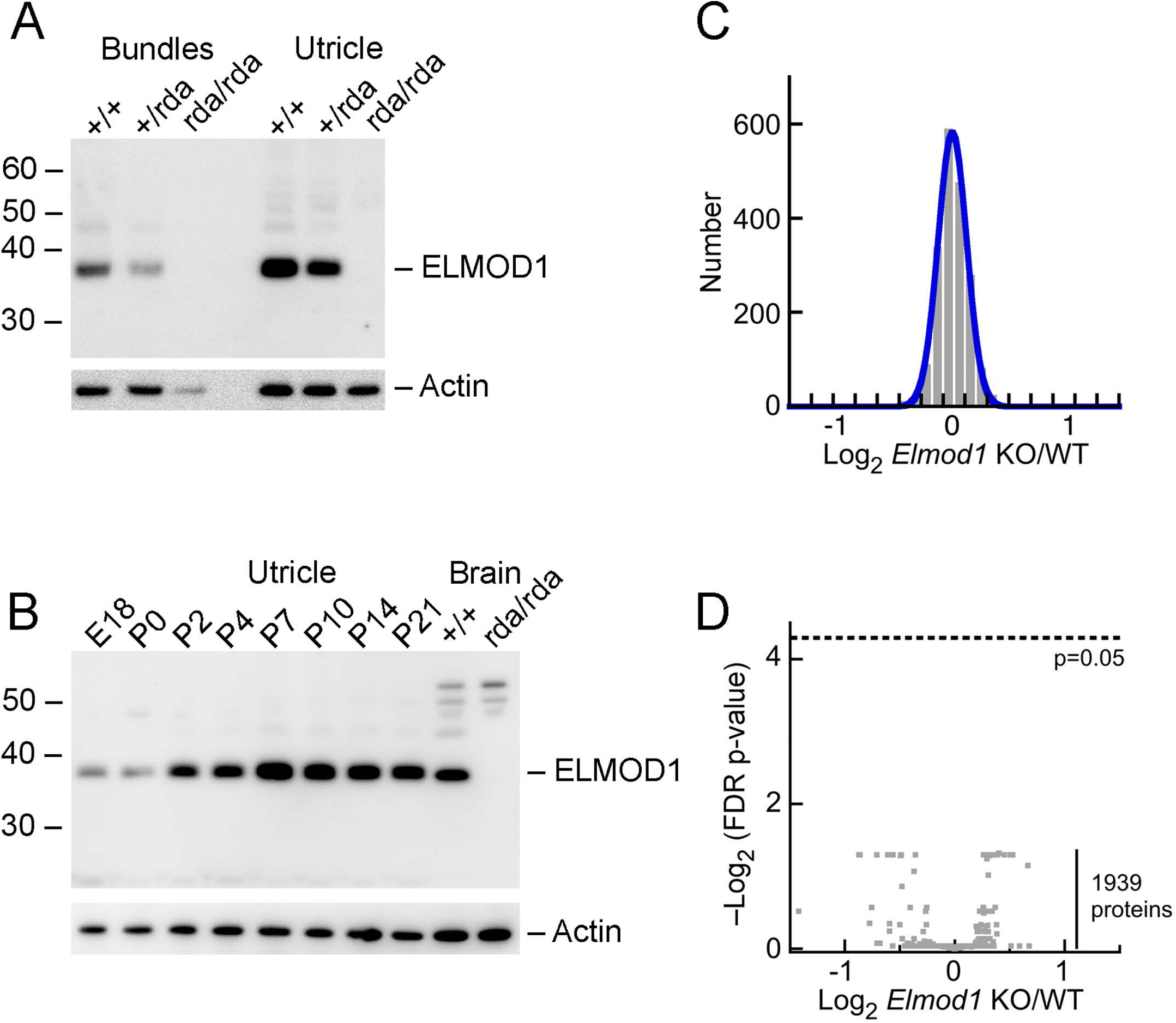
ELMOD1 protein analysis. ***A***, Immunoblot detection of ELMOD1 in mouse hair bundles and whole utricles from wild-type, heterozygous, and homozygous *Elmod1*^*rda*^ mice. Identical samples were analyzed for actin. ***B***, Developmental time course of ELMOD1 expression in whole utricle. ***C***, Log_2_ ratio of expression for 1939 proteins detected in 4/4 *rda/rda* utricle samples and 4/4 +/+ samples. ***D***, Volcano plot comparing *rda/rda* to +/+ ratio (x-axis) with FDR-adjusted p-value (y-axis). No proteins approach the p=0.05 significance level.

Because loss of ELMOD1 might lead to changes in protein content detectable by mass spectrometry, we used multiplex quantitation with isobaric chemical tags (Thompson et al., 2003) to analyze quantitatively the ~2000 most abundant proteins of the utricle and determine whether there were any significant changes in protein content due to the *rda* mutation. Proteins were isolated from *rda/rda* or *rda/*+utricles and digested with trypsin; peptides were then labeled with tandem mass tag (TMT) isobaric reagents (Thompson et al., 2003), subjected to mass spectrometry using an instrument with an Orbitrap detector, and quantified by comparison of tag ion intensities. Proteins with two or more independent peptides that were detected in both samples showed little change between the two genotypes (Fig. 2C); indeed, a volcano plot showed that none of the differences were significant (Fig. 2D). Multiple ARF family members (ARF1, ARF4, ARF5, ARF6, ARL1, ARL3, ARL8A, and ARL8B) were detected in these experiments, but ELMOD1 was not, consistent with its low expression level. Mutation of *Elmod1* thus does not lead to large-scale changes in the utricle proteome.

### Localization of ELMOD1

We localized ELMOD1 in P21 utricles, where it was enriched in hair cells with a substantial concentration in the soma’s apex (Fig. 3A-B). This apical labeling was punctate, reminiscent of intracellular membranes (Fig. 3B). No ELMOD1 signal was observed in *rda/rda* utricles (Fig. 3C-D). The phenotype originally reported for *rda/rda* cochlear hair cells, where stereocilia were fused and elongated (Johnson et al., 2012), was also seen in *rda/rda* utricle hair cells (Fig. 3C). Similar ELMOD1 localization results were seen with utricles from younger animals, but nonspecific binding of the antibody was more prominent then.

**Figure 3.**
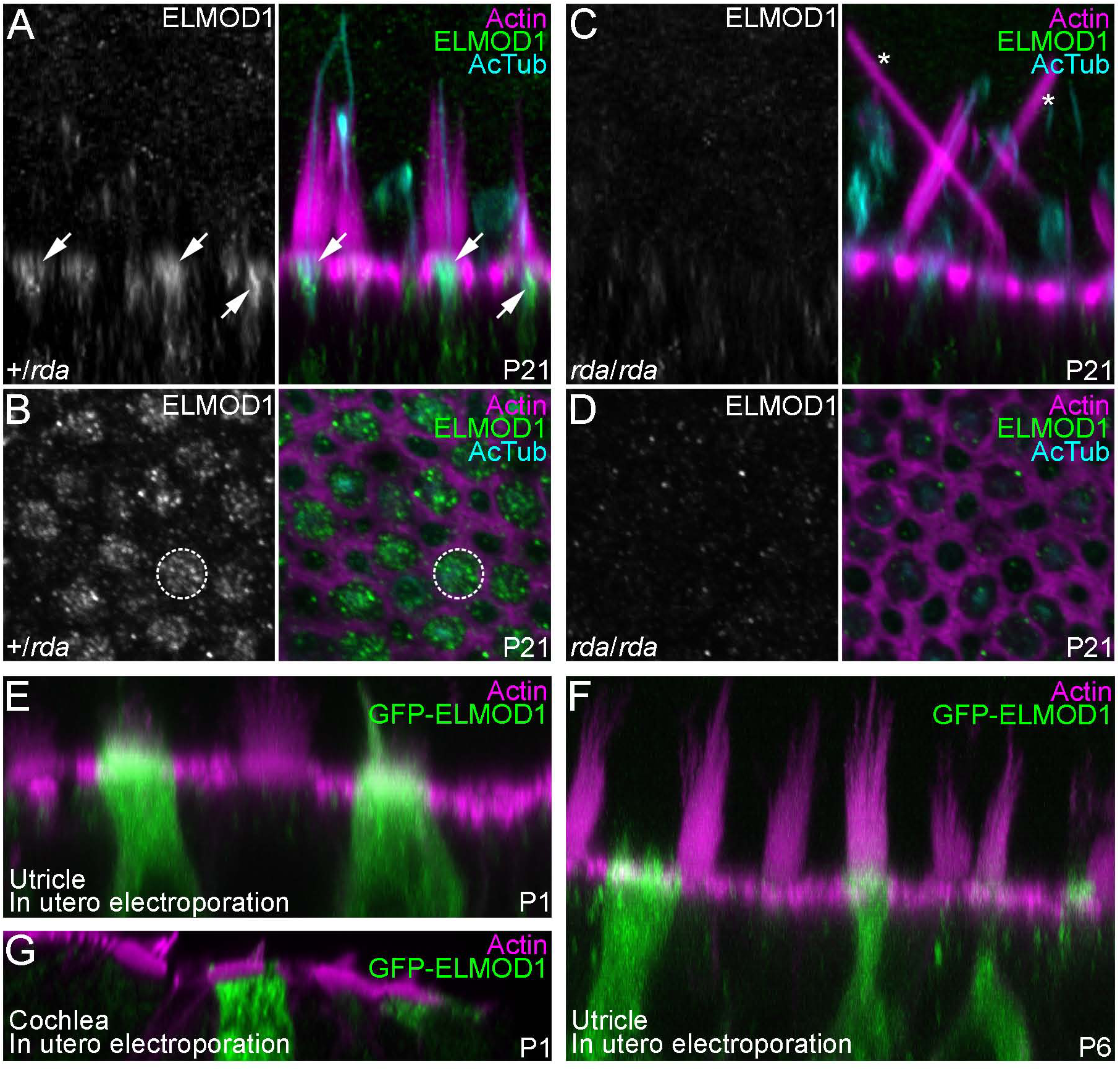
Localization of ELMOD1 in utricle hair cells. ***A-D***, Confocal images of utricle hair cells from P21 +*/rda* (A,B) or *rda/rda* (C,D) mice. Images were x-z reslices (A,C) to show ELMOD1 distribution at the apical region of hair cells (arrows) or x-y confocal sections (B,D) to show ELMOD1 at the cuticular plate level (dashed circle). No ELMOD1 signal is seen in *rda/rda* utricles (C-D); asterisks indicate giant fused stereocilia. Utricles were labeled with phalloidin (magenta) and stained with anti-acetylated tubulin (cyan) and anti-ELMOD1 (green) antibodies. ***E-G***, Confocal images of vestibular (E, F) or cochlear (G) hair bundles from mice electroporated in utero with a GFP-ELMOD1 construct at E11.5 and dissected at P1 (E, G) or P6 (F). Utricles and cochleae were labeled with phalloidin (magenta) and stained with antiGFP antibody (green) to amplify GFP-ELMOD1 fluorescence. Panel full widths: A-D, 25 μm; E, 40 μm; F-G, 50 μm.

We used in utero electroporation of GFP-ELMOD1, detected with anti-GFP for increased sensitivity, to see where exogenously expressed ELMOD1 localized in hair cells. When GFP-ELMOD1 was electroporated into E11.5 otocysts and examined at P1 (Fig. 3E) or P6 (Fig. 3F), GFP-ELMOD1 localized to the hair bundle, kinocilium, and cell body of utricle hair cells, but was particularly enriched at the apex of the cell, in the region of the cuticular plate. We occasionally detected GFP-ELMOD1 expression in cochlear hair cells. To provide higher-resolution localization of GFP-ELMOD1 there, we used image-scanning microscopy with an Airyscan detector (Müller and Enderlein, 2010; Sheppard et al., 2013; Roth et al., 2016). Some anti-GFP signal was detected in the stereocilia membrane of outer hair cells, but the anti-GFP signal was much brighter in tubular structures in the cells’ somas (Fig. 3G).

### Degeneration of hair bundles and cuticular plates in *Elmod1*^*rda*^ utricles

Actin structures of hair bundles and apical surfaces of P2 *rda/rda* utricles appeared to be very similar to those of heterozygote controls (Fig. 4). Although *rda/rda* stereocilia appeared normal at P5 and P7 (Fig. 4A), hair cells’ apical surfaces protruded modestly and gaps were present within the actin mesh of the cuticular plate (Fig. 4B, asterisks). These gaps in the cuticular plate were larger in P12 and P21 utricle hair cells (Fig. 4B), and were also detected in IHCs from P12 *rda/rda* mice (not shown). Utricles from P12 and P21 *rda/rda* mice contained fused and elongated stereocilia (Fig. 4A, arrowheads), similar to those seen in IHCs from *rda/rda* mice.

**Figure 4.**
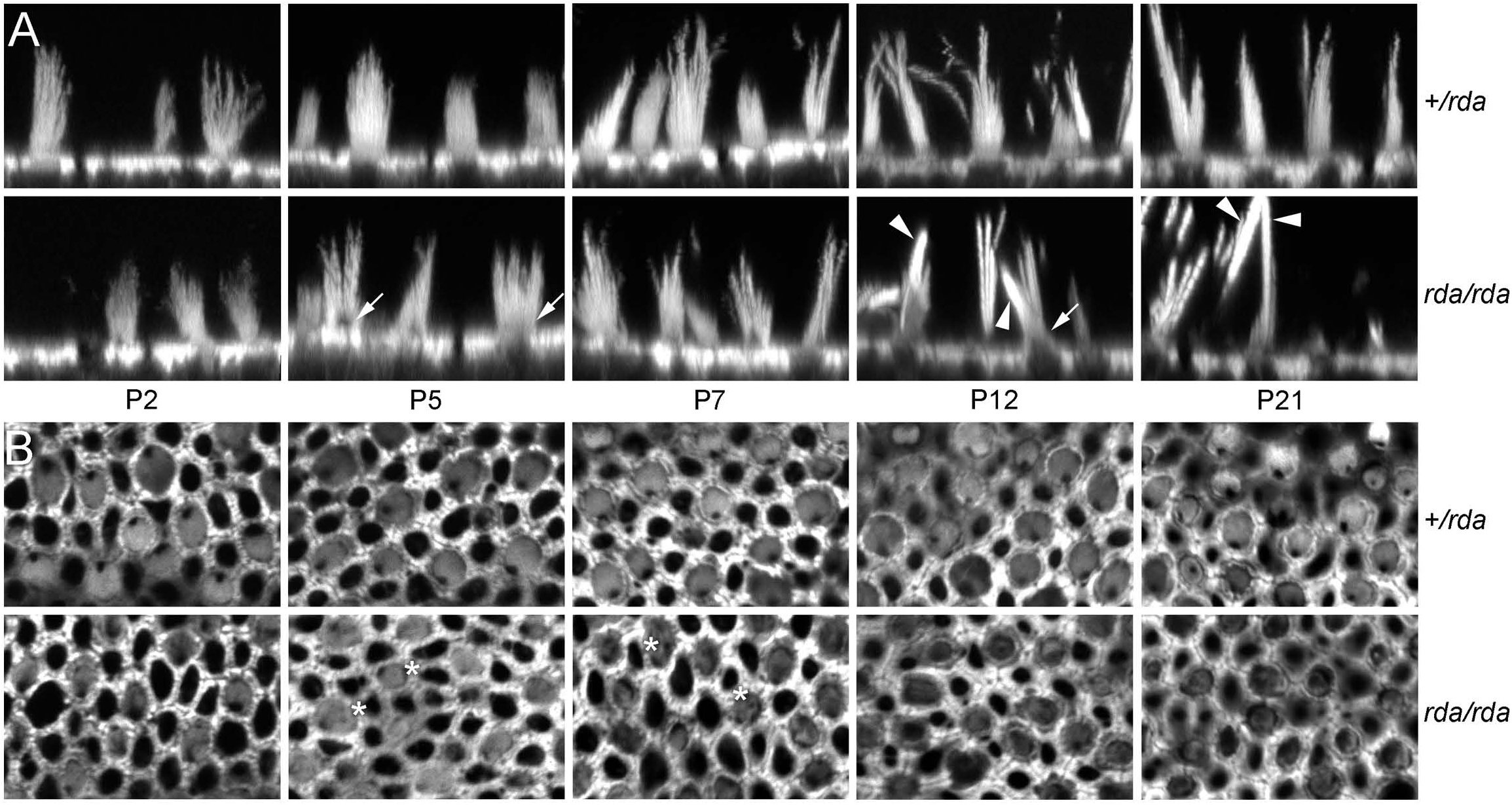
Progressive actin structural defects in *rda/rda* vestibular hair cells. ***A***, Confocal images of phalloidin-labeled vestibular hair bundles from +*/rda* (top) or *rda/rda* (bottom) mice at various postnatal ages. Arrows mark protrusions of the bundle from the apical surface (beginning at P5) and arrowheads mark fused stereocilia (beginning at P12). ***B***, Cuticular plates of vestibular hair cells degrade between P5 and P21 in *rda/rda* mice. Confocal images at the level of the cuticular plate of phalloidin-labeled vestibular hair cells from +*/rda* (top) or *rda/rda* (bottom) mice at various postnatal ages. Asterisks mark holes within the cuticular plate actin that develop starting at P5, indicating that cuticular plate degradation precedes stereocilia fusion in *rda/rda* mice. All panels are 30 μM wide.

Transmission electron microscopy (TEM) imaging of P5 utricles, prior to stereocilia fusion, showed that the number and density of vesicles and endosome-like structures below the hair bundle increased substantially in *rda/rda* mice (Fig. 5A-D); the cuticular plate appeared to be porous or missing, and many membranous structures were present where the cuticular plate normally sits (Fig. 5B', D'). Apical membranes had lifted substantially above the level of the tight junctions in *rda/rda* hair cells. We defined an angle θ that is defined by the elevation of the middle of apical surface above a line drawn between the tight junctions on either side of the cell (Fig. 5E); θ was more than 3-fold larger in *rda/rda* hair cells as compared to heterozygote controls (Fig. 5F; p<0.0001 by Student's t-test). We also noted that clathrin-coated pits and vesicles were found more frequently in *rda/rda* than in heterozygous hair cells (Fig. 5G-H).

**Figure 5.**
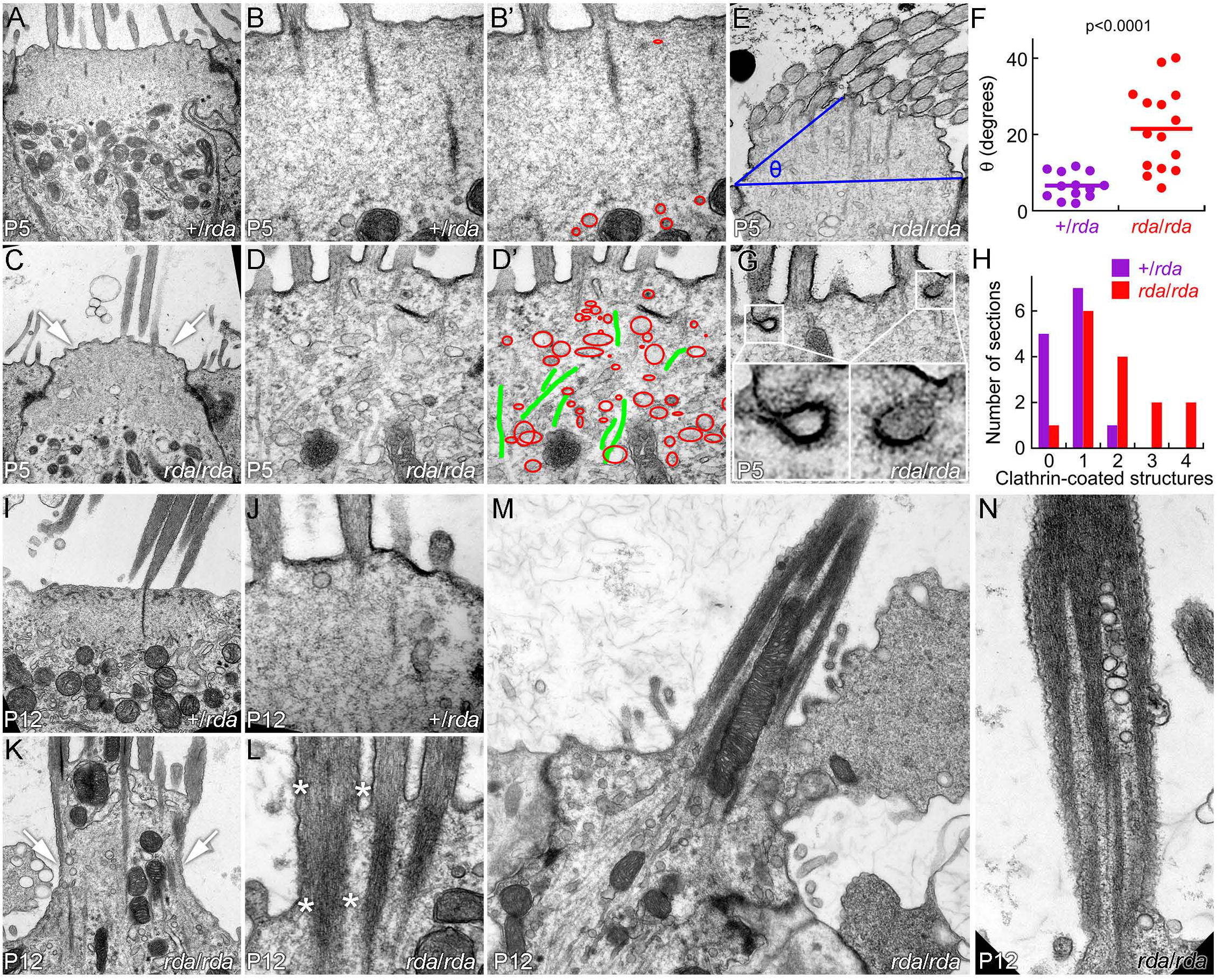
Accumulation of apical vesicles and progressive stereocilia fusion in *rda/rda* vestibular hair cells. ***A-B***, +*/rda* vestibular hair cells at P5. ***C-D***, *rda/rda* vestibular hair cells at P5. Notice bulging of apical membrane (arrows) and penetration of vesicular structures into the cuticular plate region (outlined in red). ***B’***, ***D’***, Vesicular structures (red) and microtubules (green) overlaid on B and D images.***E***, Definition of angle θ. A line was drawn connecting the apical end of the tight junctions on either side of the cell, and a second line was drawn from one tight junction to the apical surface halfway across the cell. θ is the angle defined by those two lines. ***F***, Increase in θ measured from *rda/rda* hair cells. ***G***, Examples of clathrin-coated pits and vesicles in *rda/rda* hair cells. ***H***, Quantitation of clathrin-coated pits and vesicles. The x-axis indicates the number of clathrin structures seen in a single section, while the y-axis indicates the number of sections with that number of clathrin structures. *I-J*, +*/rda* vestibular hair cells at P12. *I-J*, P12 +*/rda* hair cells. *K-N*, P12 *rda/rda* vestibular hair cells. Notice massive extrusion of the cell’s apical domain (arrows in K; see also extrusion of apical membrane in M) and fusion of stereocilia actin cores (asterisks in L; see also N). Panel full widths: A, C, I, K, 4 μm; B, D, J, L, 1.2 μm; E, 2.5 μm; G, 1.2 μm; M, 5 μm; N, 1.5 μm.

By P12, utricles from *rda/rda* mice had extensive degradation of the cuticular plate and protrusion of microtubules, mitochondria, and vesicles up towards the apical surface (Fig. 5I-L). The apical membrane in many cases blebbed outwards in large structures (Fig. 5M). Stereocilia appeared to be partially swallowed by the expanding membrane and had fused, either just at their bases (Fig. 5K-L) or throughout their length (Fig. 5M-N). Mitochondria and vesicles were also found within the fused stereocilia (Fig. 5M-N).

### Expression of ARF6, PTPRQ, and MYO6 in *rda* utricles

Mutations in *Myo6*, *Ptprq*, and *Rdx* each produce a phenotype that includes apical membrane lifting and stereocilia fusion (Self et al., 1999; Kitajiri et al., 2004; Goodyear et al., 2012), which is similar to the phenotype observed in *rda/rda* mice. MYO6 and PTPRQ were both mildly mislocalized in *rda/rda* mice (Fig. 6A-D), although much of the altered distribution appeared to be due to the protrusion of the apical membrane up along stereocilia actin cores. The extent of PTPRQ labeling on stereocilia membranes was increased somewhat towards stereocilia tips (Fig. 6D). Cell-to-cell RDX labeling was variable in *rda/rda* utricles (Fig. 6F), but did not differ significantly from that of heterozygote controls (Fig. 6E). Note that in Fig. 6F we captured several fused stereocilia cores of *rda/rda* hair cells standing vertically, demonstrating the extreme morphology of the homozygous mutant.

**Figure 6.**
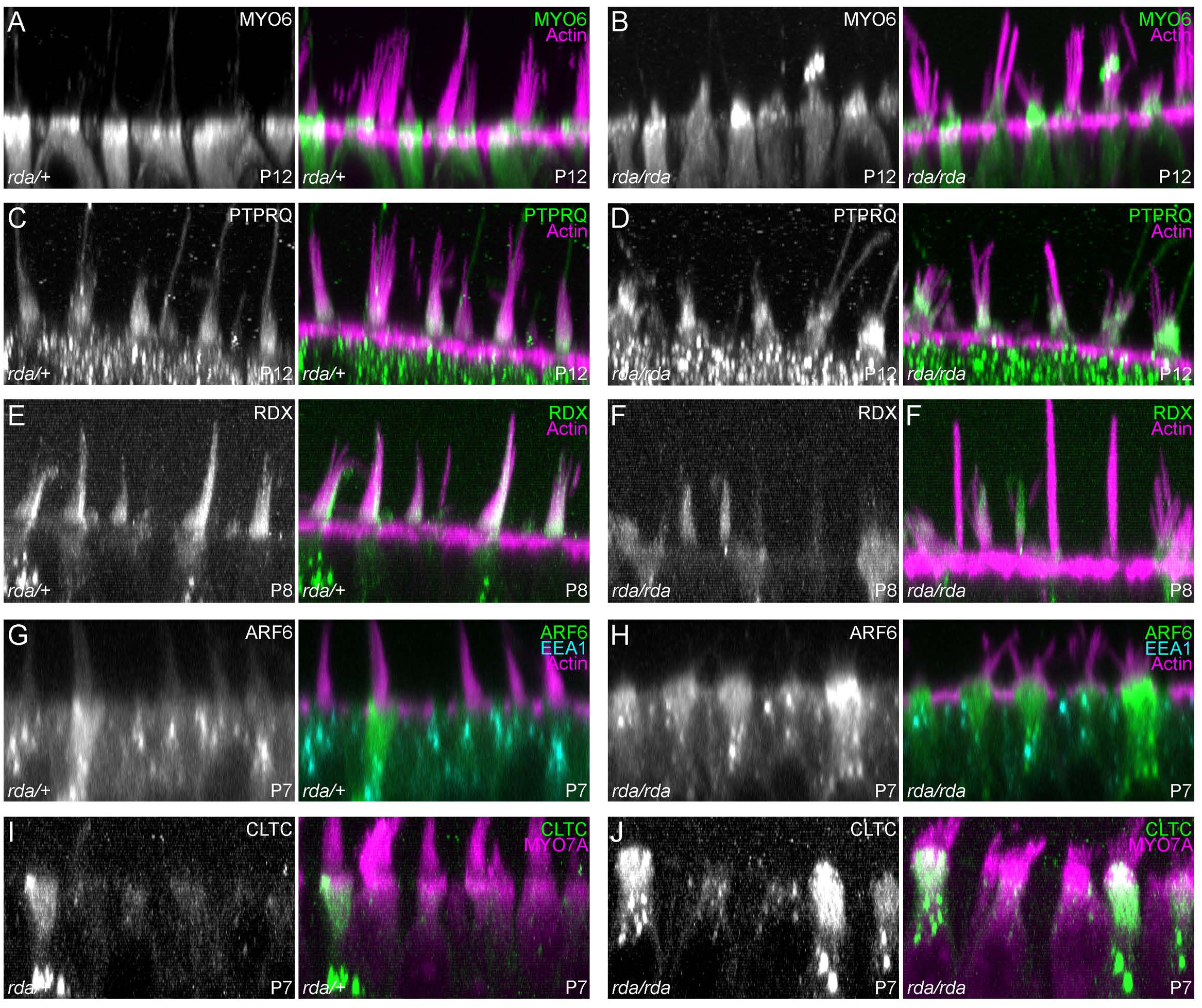
Localization of proteins in *rda/rda* vestibular hair cells. Reslice projections of confocal z-stacks of phalloidin‐ and antibody labeled vestibular hair cells from +*/rda* (left) or *rda/rda* (right) mice. ***A-B***, MYO6 in P12 utricle hair cells; maximum projection of 5 μM depth. MYO6 is present in the apical protrusions of *rda/rda* hair cells. ***C-D***, PTPRQ in P12 utricle hair cells; maximum projection of 5 μM depth. PTPRQ is also present in the apical protrusions of *rda/rda* hair cells, and in an expanded zone in hair bundles. ***E-F***, RDX in P8 utricle hair cells; average projection of 8 μM depth. ARF6 is more apically located in *rda/rda* hair cells. ***G-H***, ARF6 and EEA1 in P7 utricle hair cells; average projection of 0.8 μM depth. ARF6 is more apically located in *rda/rda* hair cells. ***I-J***, CLTC in P7 utricle hair cells; average projection of 8 μM depth. Increased number of high-expressing cells in *rda/rda* utricles. Panel full widths: 40 μm.

Because ELMOD1 is reported to be an ARF6 GAP, we also examined ARF6 labeling in *rda/rda* mice. In P7 control hair cells, ARF6 was broadly located in the apical half of hair cells, with some dense structures in supranuclear areas (Fig. 6G); by contrast, in *rda/rda* mice, ARF6 was notably focused at the apical end of many P7 hair cells (Fig. 6H). By contrast, the distribution of EEA1, an early endosome marker, was unchanged between heterozygous and homozygous mice (Fig. 6G-H). The frequency of cells expressing high levels of clathrin heavy chain increased in *rda/rda* mice (Fig. 6I-J).

### Membrane trafficking in *rda* mice

We used FM1-43 labeling to monitor membrane trafficking in hair cells (Fig. 7). When transduction-channel blockers are included in the bathing solution, FM dyes enter cells via endocytosis rather than transduction (Griesinger et al., 2002; Meyers et al., 2003). We incubated dissected utricles with the fixable dye FM1-43FX for 1 minute at 4°C, then transferred the organs to a dye-free solution at 37°C. Because the utricle’s apical membrane is exposed to extracellular solutions while the basolateral membrane is protected by apical tight junctions and the basal lamina, most dye labeling occurs through apical endocytosis, as demonstrated by the measured labeling profiles. To examine membrane trafficking in hair cells, utricles were fixed at various times after the initial pulse of dye; the resulting distribution of dye reflected membrane trafficking in the cell. In this assay, hair cells were much more robustly labeled than were supporting cells; dye was mostly located above the nucleus at 5 min post-labeling, while by 50 min there was considerable dye below the nucleus, in the synaptic region of the cell (Fig. 7A-F).

**Figure 7.**
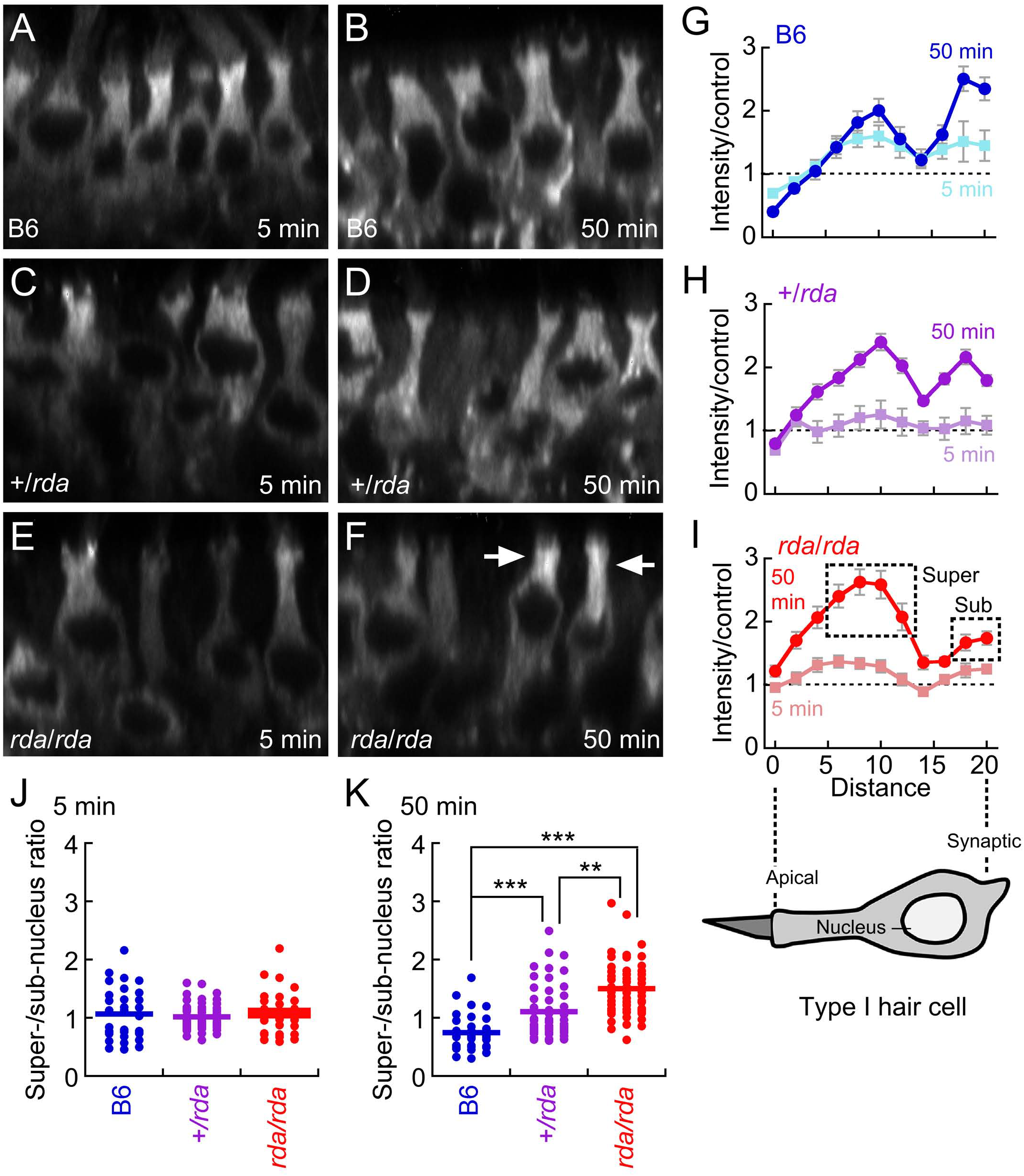
Membrane trafficking in *rda/rda* vestibular hair cells assayed with FM 1-43 labeling. ***A-F***, FM 1-43 labeling of P8 C57BL/6, *+/rda*, and *rda/rda* vestibular hair cells at 5 and 50 minutes. Each panel is an x-z reslice from a confocal stack. Arrows in F show accumulation of FM1-43 in pre-nuclear regions in *rda/rda* mutants at 50 min. All panels are 40 μM wide. ***G-I***, Quantitation of FM1-43 labeling in type I vestibular hair cells. Diagram of hair cell shows approximate location of each point along the cell. Labeling was carried out for 1 min at 4°C, then samples were shifted to 37°C for the indicated times. Fluorescence intensity profiles are normalized to profiles from samples that were labeled for 1 min at 4°C, then fixed. Points quantified for J and K are indicated by boxes in I. ***J***, Ratio of super-nucleus to sub-nucleus staining at 5 min. ***K***, Ratio of super-nucleus to sub-nucleus staining at 50 min. ***, p<0.001; **, p<0.01.

The extent of FM1-43FX dye transport within hair cells was diminished in *rda/rda* mice. We analyzed the distribution of dye over time in type I hair cells, recognized by their narrow neck above the nucleus and below the cuticular plate. We found that in comparison to B6 and +*/rda* controls (Fig. 7G-I), levels of FM1-43 in the synaptic region were reduced in *rda/rda* mice, and levels were elevated above the nucleus (Fig. 7F, arrows). We compared intensities in apical regions of the cell, above the nucleus (‘super-nucleus) to synaptic regions (‘sub-nucleus). While the genotypes did not vary at 5 min (Fig. 7J), the super-/sub-nuclear ratio was significantly elevated in *rda/rda* hair cells compared to +*/rda* or B6 controls (Fig. 7K). We noted that this ratio was also significantly elevated in +*/rda* as compared to B6 controls (Fig. 7K).

### ARF6-GTP labeling peaks at P8 in developing utricles

ARF6 protein levels were relatively constant over utricle development as shown by immunoblotting (Fig. 8A) and immunocytochemistry (Fig. 8B-D,H-J). ARF6 was significantly elevated at the cuticular plate level at P8, however, as compared to P5 (Fig. 8H-L,N). We also examined the distribution of activated ARF6 using an ARF6-GTP antibody (Torii et al., 2014; Reviriego-Mendoza and Santy, 2015). Levels of ARF6-GTP peaked around P8 (Fig. 8E-G,K-M,O), which is close to the peak of ELMOD1 expression (Fig. 2). ARF6-GTP was present at basolateral membranes in most cells (e.g., Fig. 8K), but many cells, particularly at P8, had elevated ARF6-GTP surrounding the cuticular plate (arrows, Fig. 8L). ARF6-GTP at the cuticular plate level was significantly elevated at P8 as compared to P12 but not P5 (Fig. 8O). Thus both ARF6 and ARF6-GTP peak at the cuticular plate level around P8, similar to the peak of ELMOD1 expression (Fig. 2B).

**Figure 8.**
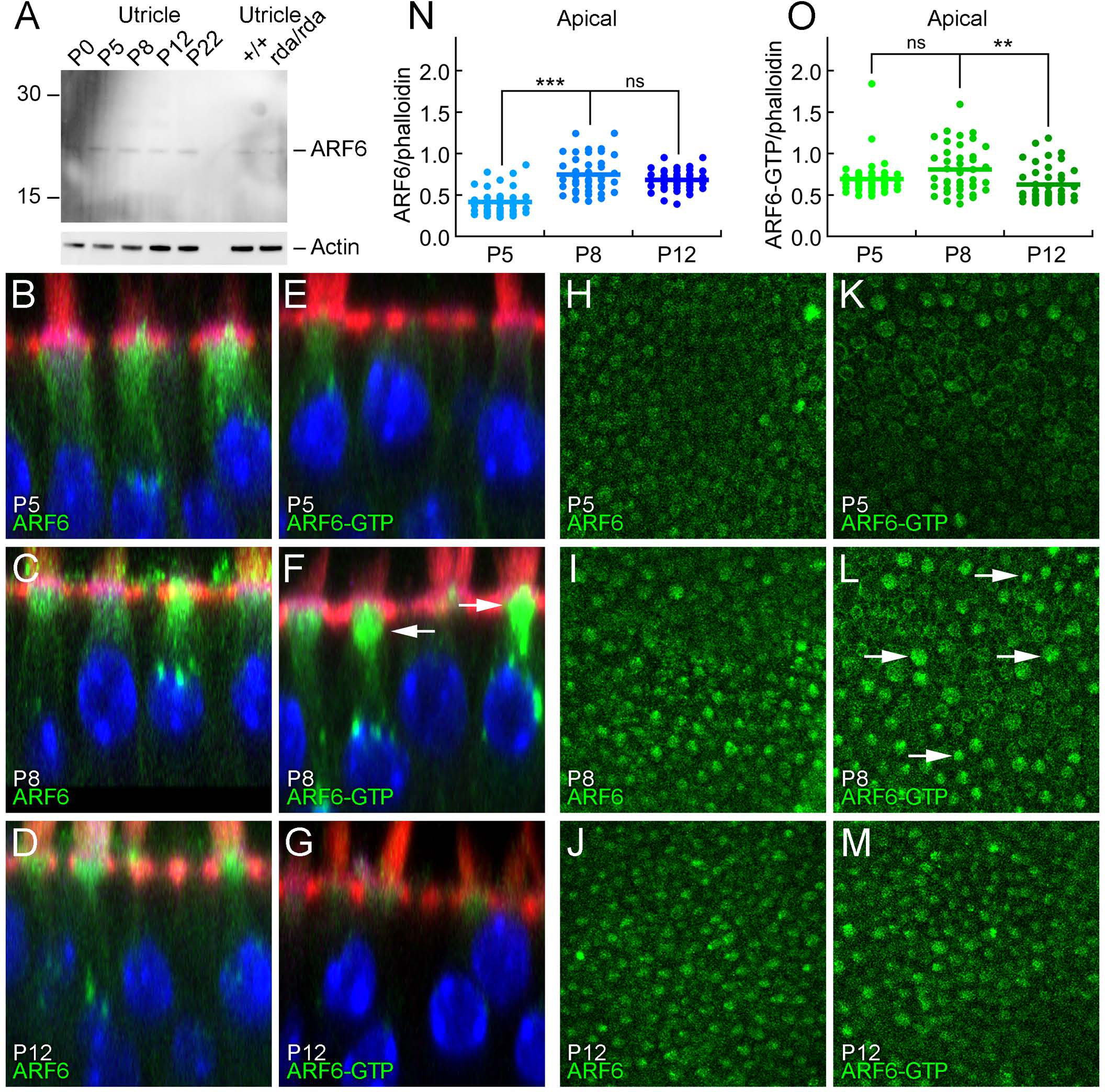
ARF6 and ARF6-GTP during hair bundle development. ***A***, Immunoblot detection of ARF6. Total ARF6 levels are constant during postnatal utricle development, and are unchanged in *rda/rda* hair cells. Blot was probed separately for actin (below). ***B-G***, ARF6 and ARF6-GTP immunolabeling in profile views of C57BL/6 utricles during development. Little change in ARF6, but ARF6-GTP is elevated in some hair cells at P8. Panel sizes: 30 × 30 pm. ***H-M***, ARF6 and ARF6-GTP immunolabeling in cross-sections of C57BL/6 utricles at the cuticular-plate level during development. Arrows indicate cells with higher levels of ARF6-GTP. Panel sizes: 100 × 100 μm. ***N***, Immunocytochemistry quantitation of ARF6 levels relative to phalloidin. ***O***, Immunocytochemistry quantitation of ARF6-GTP levels relative to phalloidin. ***, p<0.001; **, p<0.01; ns, not significant.

### Trafficking proteins in hair cells

Given the importance of ARF6 in apical membrane recycling and the possible role of ELMOD1 in regulating ARF6, we investigated in hair cells several other components of recycling pathways for apical membranes (Fig. 9). As previously noted, ARF6 was focused in the apical half of the hair cell, with some concentration in bright spots just above the nucleus (Fig. 9A,G). Consistent with ARF6’s role in clathrin-mediated endocytosis (Doherty and McMahon, 2009), the clathrin subunit CLTC had a very similar pattern to that of ARF6, with apical concentration and supranuclear aggregates (Fig. 9B,H). Aggregates containing the early endosome marker EEA1 were more numerous than those labeled for ARF6; we also noted that EEA1 staining was considerably deeper in type I hair cells as compared with type II, consistent with their deeper nuclei (Fig. 9C,I). LAMP1, which marks lysosomes, had a similar distribution to ARF6 in type I but not type II cells, where there were fewer large aggregates. RAB11, a marker of apical recycling endosomes, and RAB5, which localizes to apical early endosomes, were more uniformly distributed in hair cells (Fig. 9E-K, F-L), unlike the distribution of ARF6. Thus ARF6 appears to be associated with a subset of apical endosomal compartments, as well as with lysosomes.

**Figure 9.**
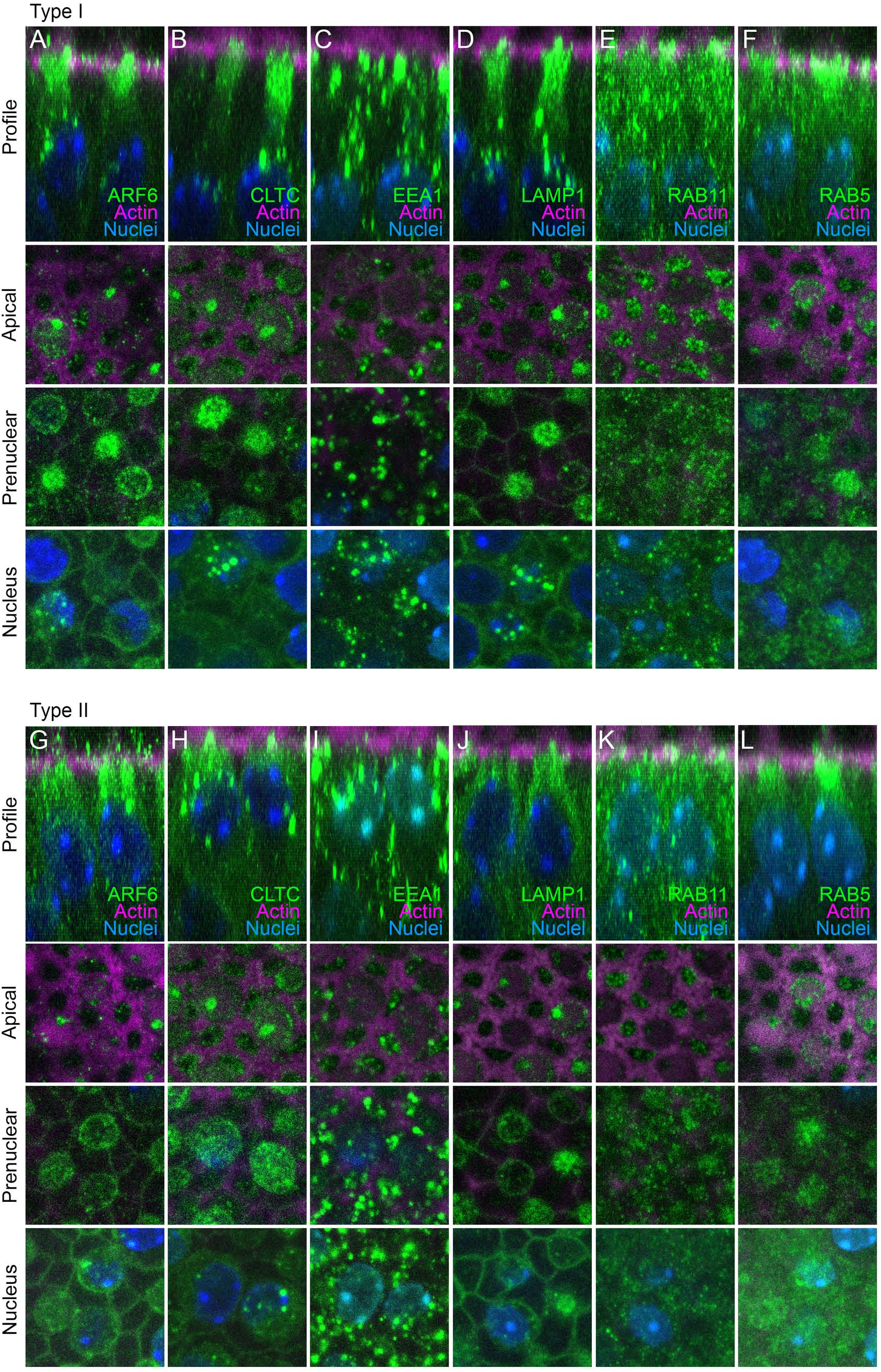
Hair-cell localization of proteins involved in apical membrane trafficking. Displayed for each protein (in type I or type II hair cells) are an x-z reslice showing the protein’s apical-basal profile, as well as x-y slices at apical, prenuclear, and nuclear levels. ***A-F***, type I hair cells. ***G-L***, type II hair cells. ***A***,***G***: ARF6. ***B***,***H***: CLTC. ***C***,***I***: EEA1. ***D***,***J***: LAMP1. ***E***,***K***: RAB11. ***F***,***L***: RAB5. Panel full widths: 18 μm.

### ARF6 GTP/GDP ratio is elevated in *Elmod1*^*rda*^ utricles

If ELMOD1 is an ARF6 GAP, the GTP/GDP ratio for ARF6 should be increased in *rda/rda* utricles. To measure ARF6-GTP levels, we carried out pulldowns using the protein binding domain (PBD) of the ARF6 effector protein GGA3, which binds GTP-bound ARF6 (Takatsu et al., 2002). We precipitated ARF6-GTP from utricle protein extracts with immobilized PBD-GGA3, then detected ARF6 with a specific antibody (Fig. 10A). Comparing precipitated ARF6 to total ARF6, we estimated that the ARF6 GTP/GDP ratio was elevated ~4-fold in P12 *rda/rda* utricles as compared to heterozygotes or B6 controls (Fig. 10B), confirming ELMOD1’s role as an ARF6 GAP in hair cells.

**Figure 10.**
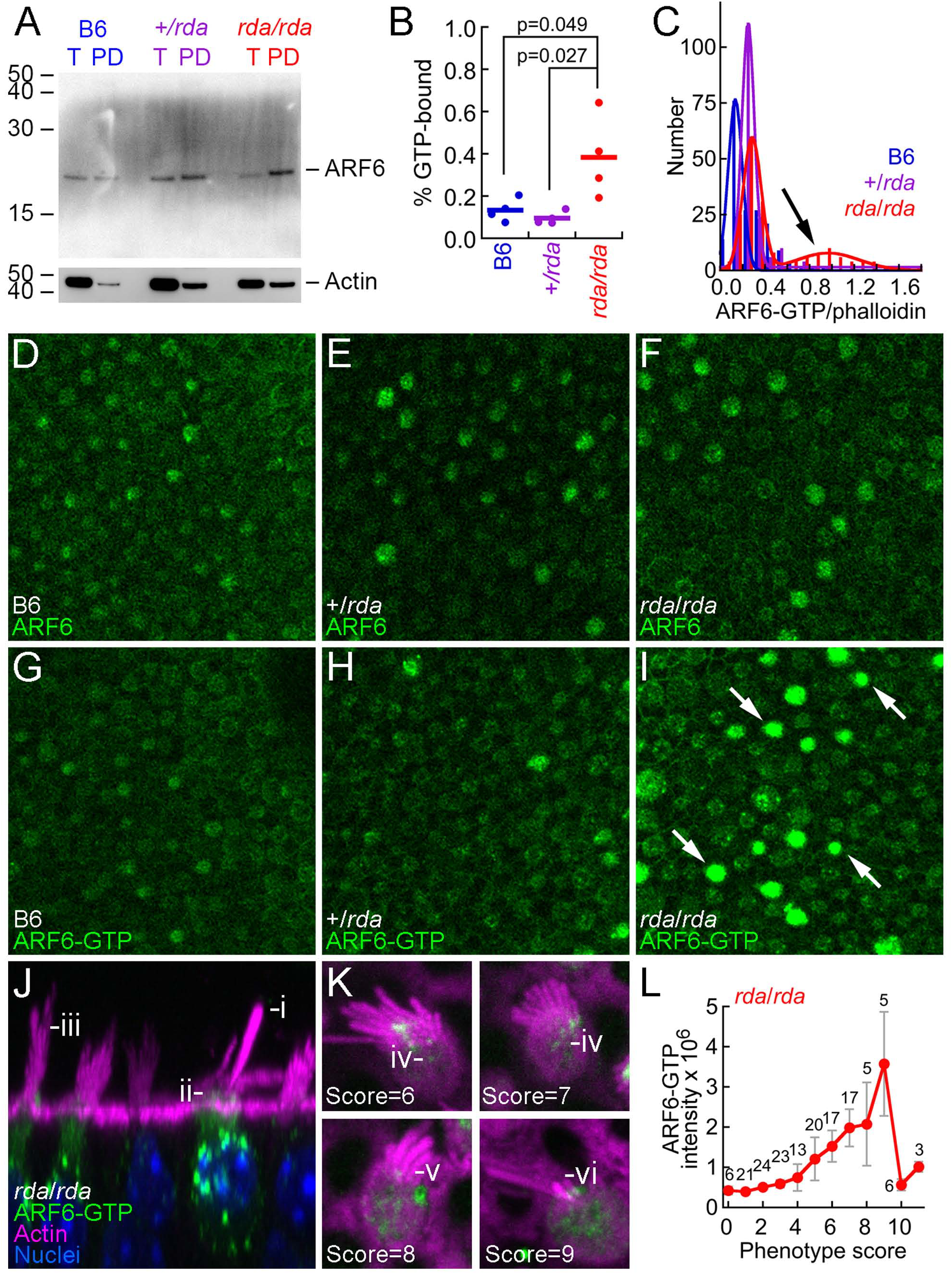
ELMOD1 is an ARF6 GAP in hair cells. ***A***, Immunoblot showing total ARF6 (T) or ARF6-GTP pulled down with PBD-GGA3 (PD) from B6, +*/rda*, and *rda/rda* utricle extracts. Blot was reprobed for actin (below). T, total material; PD, pulldown. Equal fractions of each sample were loaded, allowing direct comparison band intensity. ***B***, Summed data from four PBD-GGA3 pulldown experiments. The fraction of ARF6 in the GTP-bound state was significantly elevated in *rda/rda* utricles as compared to either +*/rda* or B6 utricles. ***C***, Quantitation of immunocytochemistry data. Two-Gaussian fits; arrow indicates population of *rda/rda* hair cells with high levels of ARF6-GTP. +*/rda* and B6 mice had smaller populations of high-expressing cells, and expression levels were not as high. ***D-I***, Examples of immunocytochemistry at cuticular-plate level of hair cells for ARF6 (D-F) or ARF6-GTP (G-I). Arrows indicate *rda/rda* cells with high levels of ARF6-GTP. Panels are 100 × 100 μm. ***J-K***, Demonstration of *rda/rda* phenotype in cells with high ARF6-GTP levels. Examples of the *rda/rda* phenotype are shown in reslice image (J) or x-y images (K). Morphology phenotypes are indicated: (i) fused and elongated stereocilium, (ii) apical protrusion, (iii) high proportion of long stereocilia in bundle, (iv) holes in cuticular plate, (v) movement of bundle to fonticulus, and (vi) reduced bundle cross-sectional area. Panels are 37 × 37 μM (J) or 10 × 10 μM (K). For a given cell, phenotype scores (0-2 scale) were summed for each of the features labeled i-vi. Total phenotype scores are indicated for each cell in K. ***L***, Summary of data. Correlation of summed phenotype score against average ARF6-GTP intensity (mean ± SEM). Number of cells with a given phenotype score are indicated above data points.

We corroborated this result using immunocytochemistry with the ARF6-GTP antibody. Total ARF6 levels were similar at the cuticular plate level in B6, +/rda, and *rda/rda* hair cells at P8, although there was substantial cell-to-cell variability (Fig. 10C-I). By contrast, the number of cells with elevated ARF6-GTP levels was notably elevated in *rda/rda* hair cells as compared to controls (Fig. 10C,I), consistent with the lack of ELMOD1 GAP activity.

### ARF6-GTP levels are correlated with the *rda/rda* phenotype

We compared the phenotypes of 160 *rda/rda* cells to their ARF6-GTP levels. Our qualitative measures of the *rda/rda* phenotype included gaps in the cuticular plate, apical protrusion, movement of the hair bundle from a central location to nearly at the fonticulus, decreased bundle cross-sectional area (perhaps due to loss of stereocilia), and stereocilia fusion (Fig. 10J-K). Each cell was graded 0, 1, or 2 for each measure, and all measures were combined to generate a phenotype score for that cell. Combining type I and II hair cells, we found that the ARF6-GTP intensity rose as the phenotype score increased (Fig. 10L), although the cells with the most disrupted morphology had low ARF6-GTP levels. We noted that while ~40% of the type I cells had high ARF6-GTP levels, ARF6-GTP was elevated in only ~20% of the type II cells.

## Discussion

We show here that ELMOD1 acts as a GTPase-activating protein for ARF6 in hair cells, and its developmentally regulated activity is required to stabilize apical structures of hair cells. The *rda* mutation, which causes the complete loss of ELMOD1 protein, leads to a hair-cell phenotype that includes fused and elongated stereocilia, degenerating cuticular plates, altered apical endocytosis, and inhibition of vesicle trafficking. These results suggest that the maintenance of both cuticular plate and stereocilia structure in hair cells requires conversion of ARF6 to the GDP-bound form, at least at the apical surface and at the appropriate time in development.

### Biasing ARF6 to the GTP-bound form underlies the *rda/rda* phenotype

ARF proteins are generally thought to be inactive in the GDP-bound state, requiring GEFs for activation and GAPs for subsequent inactivation (Gillingham and Munro, 2007). The cytohesin and PSD (also known as EFA6) families of ARF6 GEFs are all expressed in hair cells, as are the ACAP, ARAP, GIT, and SMAP GAP families. ARFGEF1-2 and KLF6 were detectable by mass spectrometry, but these ARF GEFs do not activate ARF6 (Yamauchi et al., 2017). Many GAPs are also expressed in inner ear tissues; ELMOD1 was by far the most enriched GAP in hair cells, however, so it may be responsible for any GTPase activation that must take place only in hair cells.

Biochemical evidence shows that ELMOD1 is an ARF6 GAP in vitro (Ivanova et al., 2014). Our ARF6-GTP pull-down experiments indicate that the loss of ELMOD1 substantially increases levels of ARF6-GTP, suggesting that ELMOD1 is the dominant ARF6 GAP in the utricle. Moreover, RNA-seq and localization experiments suggest that ELMOD1 is substantially enriched in hair cells over supporting cells in the utricle, which we confirmed using immunocytochemistry. Although utricle hair cells are born over several weeks, the rate of hair cell production peaks around P0 (Burns et al., 2012). Peak expression of ELMOD1 at P7 suggests that it is not needed for early steps in apical development in hair cells, but rather that its activity is needed later in development to stabilize hair bundles, cuticular plates, and apical membranes. The timing of ELMOD1 expression is also consistent with the onset of morphological changes in *rda/rda* mice, which starts at P5.

ARF6 also concentrates at cuticular plates between P5 and P12, with ARF6-GTP levels peaking at P8 in wild-type utricles. In the absence of ELMOD1, elevated ARF6-GTP levels are seen in many P8 cells. Moreover, elevated ARF6-GTP levels are correlated with the *rda* phenotype, which suggests that ELMOD1 prevents ARF6 activation in the apical domain and stabilizes actin structures during late postnatal development. Taken together, these observations suggest that ARF6-GTP must be converted to the GDP-bound form only after initial stages of hair-bundle development. Given that ELMOD1’s only known biological activity is its ARF6 GAP activity, we suggest that the phenotype seen in the *rda/rda* mice is due to the increased ARF6-GTP levels present, and that any ELMOD1 biochemical or structural effects in hair cells are mediated through ARF6.

### Inactivation of ARF6 at apical surfaces during hair-cell development

ARF6 has wide-ranging effects on vesicle trafficking, apical membrane function, and actin cytoskeleton structure that vary in cell type, presumably due to differences in expression of ARF GEFs and GAPs (D’Souza-Schorey and Chavrier, 2006; Donaldson and Jackson, 2011). Particularly relevant for hair cells are ARF6’s modulation of vesicle-recycling pathways (D’Souza-Schorey and Chavrier, 2006) and regulation of the actin cytoskeleton through PI(4,5)P_2_ production and RHO-family GTPase activation (Myers and Casanova, 2008). ARF6 activation stimulates both clathrin-mediated and clathrin-independent endocytosis; inactivation of ARF6 is then necessary for further trafficking to recycling endosomes (Naslavsky et al., 2003) and for basolateral sorting of vesicles from the recycling endosomes cells (Shteyn et al., 2011). Recycling from the tubular endosomal membrane to the plasma membrane also requires ARF6 activation (Radhakrishna and Donaldson, 1997). ARF6 activation also leads to increased Rac1 activity and the production of PI(4,5)P2, both of which are tied to its regulation of endocytosis and endosomal trafficking (Brown et al., 2001; Palacios et al., 2002; Donaldson, 2005).

Because ELMOD1’s absence in *rda* mutants leads to elevated ARF6-GTP levels, the *rda* phenotype may be similar to phenotypes of cells expressing mutant ARF6 molecules that are constitutively active, such as ARF6Q67L. ARF6Q67L stimulates clathrin-mediated endocytosis at the apical surface of polarized epithelial cells (Altschuler et al., 1999), consistent with the increase in clathrin structures in *rda/rda* hair cells. Expression of ARF6Q67L also leads to the accumulation of endocytosed vesicles in large vacuole-like structures coated with PI(4,5)P_2_ and actin, as well as inhibition of trafficking to late endosomes and subsequent recycling (Brown et al., 2001; Naslavsky et al., 2003; Naslavsky et al., 2004). ARF6Q67L expression also prevents basolateral targeting of endosomes in polarized epithelial cells (Shteyn et al., 2011).

Many features of the *rda/rda* hair-cell phenotype are consistent with the proposed role of ARF6 inactivation in endocytic trafficking. In the absence of ARF6 inactivation by ELMOD1, endocytosed vesicles accumulated at the apical surface of P5-P8 hair cells and basolateral trafficking of FM1-43 labeled vesicles was reduced. Our results suggest that as the hair bundle matures, there may be an increased demand for apical membrane trafficking that requires ARF6 activation; ELMOD1 upregulation may be needed then to inactivate ARF6, allowing for proper trafficking and sorting of endocytosed vesicles. In the absence of ELMOD1-mediated ARF6 inactivation during this time period, excessive levels of ARF6-GTP lead to the apical accumulation of endocytosed vesicles and the decrease in apical plasma membrane seen in *rda/rda* mutant bundles.

The progression of the phenotype within *rda/rda* vestibular hair cells suggests that misregulation of endocytic trafficking and cuticular plate actin may drive the subsequent changes in hair-bundle morphology. Prior to any stereocilia changes, the apical membrane lifts, gaps appear in cuticular plate actin, endocytosed vesicles accumulate within the cuticular plate. As cuticular-plate gaps increased, apical surfaces protrude and the bundle moves towards the fonticulus, the hole in the cuticular plate through which the kinocilia inserts. Simultaneously, the cross-sectional area of the bundle decreases, perhaps as stereocilia disappear, and the remaining stereocilia increase in length and fuse. Stereocilia fusion in *rda/rda* mutants may result both from the decrease of membrane tethering to the cuticular plate as well as a decrease in the total amount of apical membrane available. Whether the gaps in cuticular-plate actin are related to the changes in endocytosis is unclear.

### *Elmod1 rda* mutant mouse shares phenotypes with *Myo6*, *Ptprq*, and Rdx mutants

Sharing of phenotypes by multiple genes raises the possibility that they operate in a molecular pathway. For a different example in hair cells, the Usher syndrome type I deafness mutants all show disorganized stereocilia, consistent with the suggestion that the Usher proteins are responsible for constructing interstereocilia linkages (Cosgrove and Zallocchi, 2014). Another common phenotype seen in hair cells is shared by *Myo6*, *Ptprq*, and *Rdx* mutants, which all have protruding apical membranes and eventual stereocilia elongation and fusion (Self et al., 1999; Kitajiri et al., 2004; Goodyear et al., 2012; Seki et al., 2017); a similar phenotype is seen with conditional *Cdc42* mutants (Ueyama et al., 2014). Utricle hair cells of *rda/rda* mice clearly share this phenotype, as do cochlear hair cells (Johnson et al., 2012). MYO6, PTPRQ, and RDX appear to be distributed relatively normally in *rda/rda* hair cells, which suggests that if the three proteins are in a common pathway, then ELMOD1 acts downstream of MYO6, PTPRQ, and RDX.

A plausible common pathway for ARF6, MYO6, PTPRQ, and RDX is regulation of or response to PI(4,5)P_2_ levels on the hair cell’s apical surface. In many cell types, ARF6-GTP stimulates production of PI(4,5)P_2_ at cell apical membranes (Donaldson and Jackson, 2011). Hair cells compartmentalize PI(4,5)P_2_ so that levels of the lipid are very low at stereocilia tapers and apical surface, in part due to concentration of the lipid phosphatase PTPRQ at those locations (Hirono et al., 2004; Zhao et al., 2012). MYO6 is thought to localize PTPRQ at the stereocilia bases (Sakaguchi et al., 2008), while RDX requires a localized PI(4,5)P_2_ signal in stereocilia (Zhao et al., 2012). MYO6 also has a high affinity binding site in its tail for PI(4,5)P_2_-containing liposomes (Spudich et al., 2007) and like ARF6, has been shown to regulate the trafficking of newly endocytosed vesicles (Aschenbrenner et al., 2003; Frank et al., 2004). Together, these observations suggest that *rda/rda* hair cells may have altered PI(4,5)P_2_ levels at apical surfaces, which may partly underlie the hair-bundle phenotype.

### Model for ELMOD1 function

During development of hair bundles, ARF6 is recruited to the hair cell’s apical region and activated, so that ARF6-GTP levels increase there to control actin structures and membrane trafficking. ELMOD1 apparently inactivates ARF6 soon after it reaches its peak, so a given cell will show only a transient rise in ARF6-GTP. In *rda/rda* mice, lacking ELMOD1, ARF6-GTP levels reach a peak and then remain high (Fig. 11B), leading to an increased number of cells with elevated ARF6-GTP levels at P8 (Fig. 10C). High levels of ARF6-GTP are correlated with the apical phenotypes seen in *rda/rda* mice (Fig. 10L), suggesting strongly that the persistence of activated ARF6 directly or indirectly produces abnormally long and fused stereocilia, as well as cuticular plate disruption. Why the hair cell activates ARF6 transiently at apical surfaces during hair bundle development is unclear, although several of the various relevant functions ascribed to ARF6—endocytosis, actin remodeling, PI(4,5)P_2_ accumulation—could be important. Nevertheless, our results suggest that the principal function of ELMOD1 is to inactivate ARF6, which allows cuticular plates and stereocilia to develop their mature structures.

**Figure 11.**
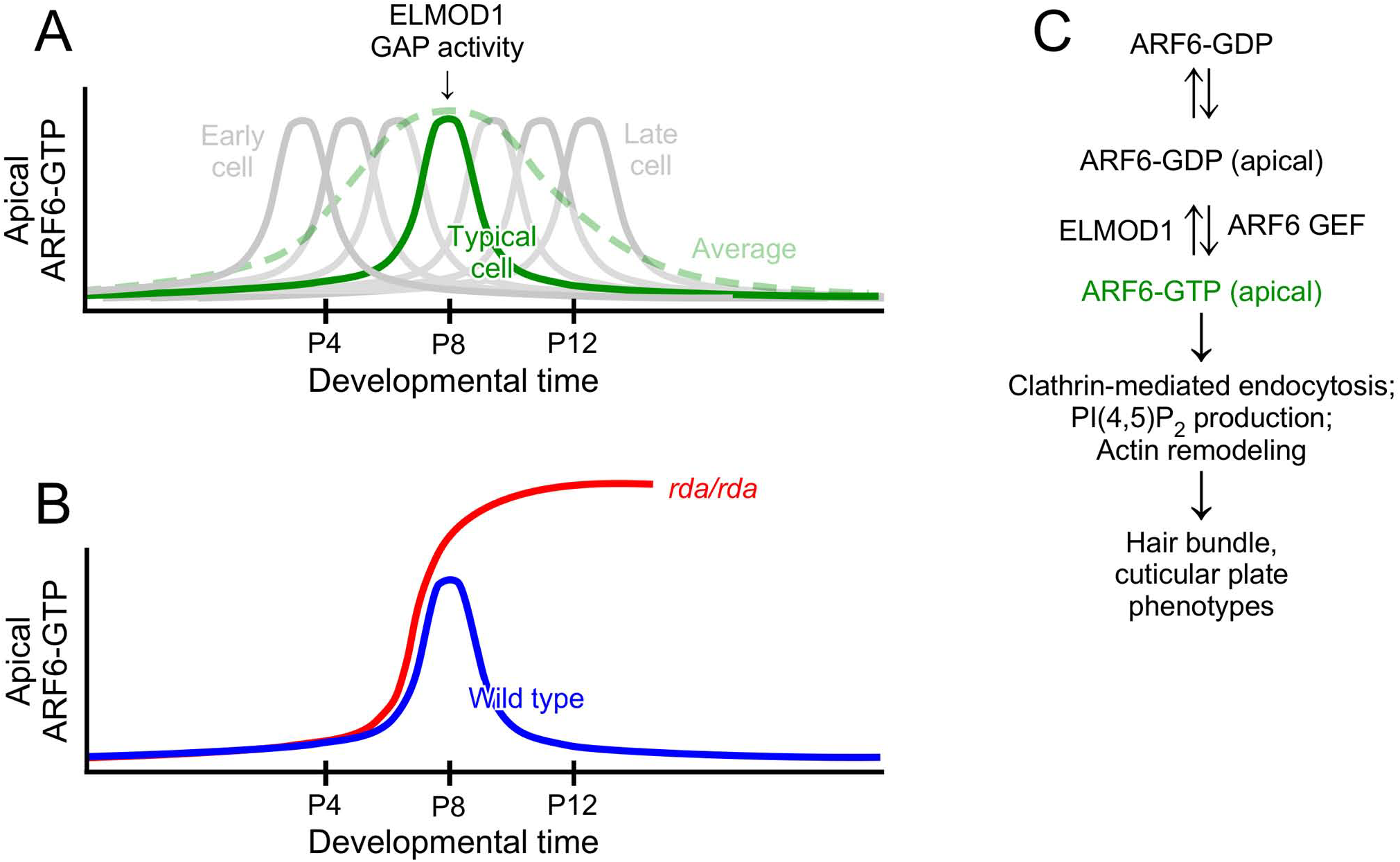
Model for inactivation of ARF6 by ELMOD1. ***A***, While ARF6-GTP (and ELMOD1) levels appear to have a broad peak during development, the average response (green dashed line) obscures the sharper peaks of individual cells. ELMOD1 levels peak around the time ARF6-GTP levels do, which produces a transient ARF6-GTP response. ***B***, In the absence of ELMOD1, ARF6-GTP levels are maintained at high levels, increasing the fraction of cells that have elevated ARF6-GTP. ***C***, Biochemical pathways for ARF6 and ELMOD1.

## Acknowledgements

We thank David Corey for sharing transcriptomics data prior to publication. PGBG was supported by NIH grants R01 DC002368 and P30 DC005983. We received support from the following core facilities: mass spectrometry from the OHSU Proteomics Shared Resource (partial support from NIH core grant P30 EY010572), confocal microscopy from the OHSU Advanced Light Microscopy Core @ The Jungers Center. Hybridoma cells for JLA20 (deposited by J. J.-C. Lin) and DSHB-GFP-4C9 were obtained from the Developmental Studies Hybridoma Bank, created by the NICHD of the NIH and maintained at The University of Iowa, Department of Biology, Iowa City, IA 52242.

We thank Ruby Larisch for mouse husbandry and Michael Bateschell for in utero electroporation. We received support from the following core facilities: mass spectrometry from the OHSU Proteomics Shared Resource (partial support from NIH core grants P30 EY010572 and P30 CA069533; Orbitrap Fusion S10 OD012246), and confocal microscopy from the OHSU Advanced Light Microscopy Core @ The Jungers Center (P30 NS061800 provided support for imaging). PGBG was supported by NIH grants R01 DC002368 and P30 DC005983. JFK was supported by NIH grant F32 DC12455.

